# Acridine Orange dye for long-term staining and live imaging of cnidarian development and regeneration

**DOI:** 10.1101/2025.11.10.687709

**Authors:** Vaidehi Patel, Valeria Dountcheva, Labib Rouhana

## Abstract

As the sister phylum to Bilateria, Cnidaria, holds a phylogenetic position that is crucial in evolutionary studies of animal development. However, the analysis of cnidarian species, as well as other emerging models, is limited by the scarcity of tools that are readily available and applied with ease. Commercial fluorescent dyes are an accessible option to increase visibility when transgenic approaches are out of reach. In this study, the commercial dye Acridine Orange (AO) was found to be suitable for long-term staining and live imaging of the cnidarian model *Nematostella vectensis.* Temporary incubation in media containing AO resulted in broad cellular labeling. Individuals stained with AO as zygotes retained the dye throughout embryogenesis, metamorphosis, and for longer than three months of growth, at which point the signal became restricted to subsets of cells. In contrast, staining of zebrafish zygotes with AO resulted in signal decay within the first couple of days of development and preferential accumulation in the yolk sac. AO is retained in *N. vectensis* adults after amputation and inherited by new tissue during regeneration. Surprisingly, AO was not found suitable for cell tracking via microinjection, as the dye dissipated within minutes of being injected into the cytoplasm of zygotes and blastomeres. Surprisingly, the subcellular distribution of AO signal did not match the pattern expected from it use in mammalian cell lines, where green fluorescence is observed when associated with cellular DNA and red fluorescence when bound to RNA. Instead, long-lived green fluorescence was observed throughout the cell surface and/or in the cytoplasm of *N. vectensis* stained with AO, whereas red emission was similarly distributed but short lived. Expanding the use of AO as a visualization tool will facilitate future studies of cnidarians and other underexplored animal models.

## INTRODUCTION

The study of species from early-diverging animal lineages provides essential insights into the evolutionary origins of developmental processes. Cnidarians, which include jellyfish, corals, and sea anemones, occupy a key phylogenetic position as the sister lineage that diverged prior to the emergence of Bilateria. In addition to their evolutionary significance, cnidarians are important in balancing ecosystems by providing homes and food for many organisms, facilitating cycling of nutrients, and by predation (Doyle et al., 2015; Boreo et al., 2005). Unfortunately, many cnidarian species, and in particular coral species, have become endangered due to rising temperatures in their habitats (Weis, 2008; Rossi et al., 2019).

Amongst cnidarians, *Hydra* have long been used in laboratories to understand cellular and molecular processes involved in development and regeneration (Trembley, 1744; reviewed by Galliot, 2012). Yet a second cnidarian species, the starlet sea anemone *Nematostella vectensis*, is gaining track as a model for investigating the evolutionary history of mechanisms involved in embryonic development (Martindale et al., 2004; Ryan et al., 2007; Finnerty et al., 2003; 2004), germline development (Chen et al., 2020; Miramón-Puértolas et al., 2024; Denner et al., 2024), and regeneration (Burton and Finnerty, 2009; Passamaneck and Martindale, 2012; DuBuc, Traylor-Knowles et al., 2014; Amiel et al., 2015; Warner et al., 2018; Cheung et al., 2025). *N. vectensis* is a diecious species for which spawning protocols of males and females are routinely performed for developmental studies under laboratory settings (Hand and Uhlinger, 1992; Stefanik et al., 2013; Layden et al., 2013; Carvalho et al., 2025). The husbandry of *N. vectensis* in the laboratory is relatively simple, as these organisms can thrive in a wide range of temperatures in natural or artificial salt water, can be maintained on a diet of brine shrimp, and as invertebrates, face fewer regulatory experimental restrictions compared to vertebrate counterparts.

Fluorescence labeling techniques enhance visualization of dynamic developmental processes in live organisms. *N. vectensis* is particularly suitable for fluorescence labeling as their external surface is largely transparent throughout much of their development and adult life. Eggs and zygotes are filled with yolk that is milky in color but becomes progressively translucent during embryonic development and metamorphosis, which take about a week. While temporary and stable transgenic expression of fluorescent proteins have been established for visualizing cellular and subcellular dynamics during development of *N. vectensis* (DuBuc, Dattoli et al., 2014; Paix et al., 2023), these are based on advanced molecular biology techniques and require equipment for microinjection. An alternative approach to fluorescent labeling is the use of commercially available dyes, some of which can be easily taken up by multiple strains, mutants, or species related to a given model. Unfortunately, the use of some commonly used dyes is limited by photobleaching, toxicity, or rapid clearance, which restrict their utility in long-term studies (Ettinger and Wittman, 2014). A suitable labeling method for live, developing cnidarians must therefore be both non-toxic and persistent over time.

In this study, we evaluated the use of a florescent dye as a long-term non-toxic tool for the visualization of early developmental processes and regeneration in *N. vectensis*. Acridine orange (AO), a fluorescent dye traditionally used for labeling nucleic acids and identifying apoptotic cells in mammalian models and cell lines (Plemel et al., 2017), was found to facilitate long-term labeling and live imaging in *N. vectensis.* AO labeling by brief immersion of embryos in media containing the dye persists from the earliest stages of development through embryogenesis, metamorphosis, and into adulthood. The dye is also retained upon amputation and inherited by regenerated tissue. While AO staining of *N. vectensis* does not follow that distribution observed in mammalian models, our results indicate that it can facilitate long-term, non-invasive, fluorescence imaging in cnidarians, therefore enabling deeper exploration of their biology.

## MATERIALS AND METHODS

### Laboratory Husbandry of N. vectensis

*Nematostella vectensis* was obtained from Mark Martindale’s laboratory at the Whitney Laboratory for Marine Bioscience (University of Florida) and cultivated at the University of Massachusetts Boston in 1/3 Artificial Salt Water (ASW). 1X ASW was made by mixing 395 g/L of Instant Ocean marine salt (Spectrum Brands, Blackburn, VA) in ultrapure (MilliQ-purified) water and diluted in ultrapure water to make 1/3 ASW. Glass bowls containing *N. vectensis* at a wide range of densities were maintained in incubators set to 16 °C and continuous dark. The animals were fed at room temperature on the benchtop twice per week with Artemia (brine shrimp) and their bowls cleaned after feeding. Sexually mature animals (≥3 months post fertilization; 3 mpf) were induced to spawn using a protocol based on Stefanik et al. (2013) that involves a meal of minced oyster flesh followed by cleaning and incubation at 26 °C under constant light for 8 hours. Husbandry media in the bowls was exchanged to 50-70% fresh husbandry media at room temperature, and bowls were left for up to 2 hours on benchtops to induce spawning. To expand the colony, eggs from spawning events were fertilized by placing them for 30 minutes in a bowl with husbandry media containing sperm from bowls with males induced to spawn concurrently. Embryos were de-jellied by being placed into a 4% cysteine solution for five minutes on a rocker before being washed with 1/3X ASW six times or more.

### Preparation and storage of Acridine Orange

To make a stock solution of 6.664 mg/mL Acridine Orange (Catalog no. A1301, Thermo Fisher Scientific, Waltham, Massachusetts), 100 mg of AO was added to 15 mL of ultrapure water, dissolved by gentle agitation, aliquoted in 1 mL volumes, and stored at -20 °C. Each aliquot was frozen and thawed a maximum of five times. Dilutions were made in ultrapure water accordingly.

### Staining of live animals with Acridine Orange

*Nematostella vectensis* embryos, polyps, and adults were incubated with 0.664 µg/mL AO in 1/3 ASW in the dark at room temperature for 45 minutes. Other concentrations were tested as indicated in (Figure 1). Samples were then washed 3-5 times with 1/3 ASW. Stained animals were maintained in 6 well plates (Product no. 31736, Costar, Arlington County, VA) alongside organisms not exposed to any dye.

**Figure 1.**
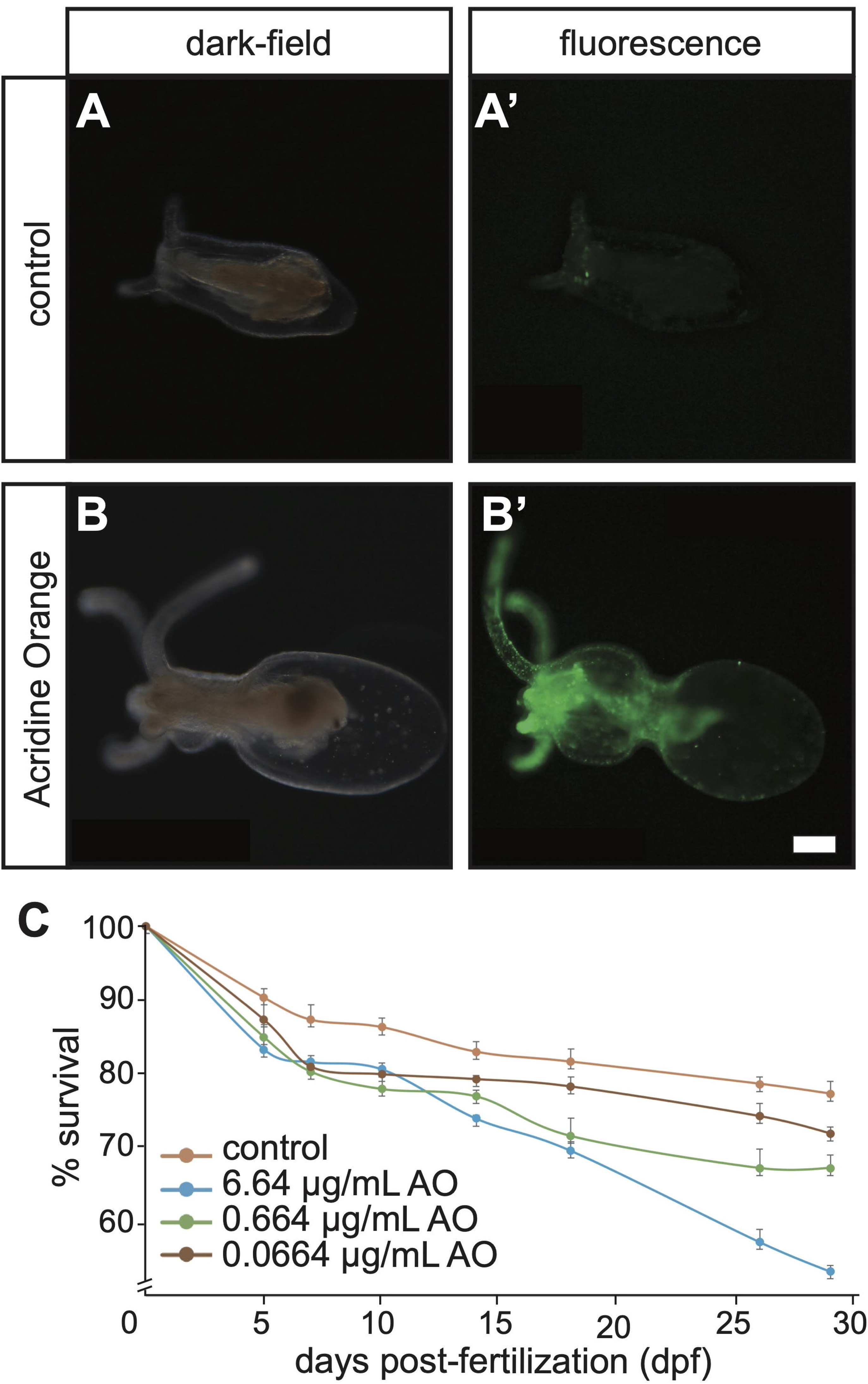
Exposure to AO allows live imaging of *N. vectensis* via green florescence microscopy. **(A-B)** *N. vectensis* polyps (46 days post-fertilization) observed under dark-field (A and B) and green fluorescence microscopy (A’ and B’) in the absence of AO exposure (A-A’) and following exposure to AO days prior (B-B’). Scale bar = 0.1 mm. **(C)** Survivability assay reveals timing and extent of mortality for *N. vectensis* embryos exposed to 0 (orange line), 6.64 (blue line), 0.664 (green line), or 0.0664 (burgundy line) µg/mL AO. Error bars at each time point represent standard deviation from the mean (n ≥ 3).

Zebrafish embryos were obtained from the Siegfried lab at University of Massachusetts Boston and maintained in standard conditions approved by the Institutional Animal Care and Use Committee (protocol #32). Zebrafish zygotes were stained by incubation in E3 media supplemented with 0.664 µg/mL of AO for 45 minutes at room temperature followed by washes with E3 media until clear. Zebrafish embryos were placed back in standard 28 °C husbandry conditions and imaged at different timepoints up to five days post-fertilization.

### Microinjection of N. vectensis blastomeres

To prepare the injection, Kwik-Fil glass capillaries (World Precision Instruments, Sarasota, FL) were pulled into needles using a P-97 Micropipette Puller (Sutter Instruments, Novato, California). The settings were adjusted to a 546 heat, 35 velocity, 95 pull, and 100 ms time to produce needles of appropriate dimensions. A solution of 66.4 µg/mL of AO in ultrapure water was loaded onto the needle and attached to a Femtojet microinjector (Eppendorf, Hamburg, Germany). *N. vectensis* zygotes and embryos that reached the 4-cell stage were injected and transferred to a new container containing 1/3 ASW solution to prevent AO leakage from staining uninjected cells. Dye maintenance and dissipation was tracked via live green florescence microscopy using a ZEISS Axio Zoom V16 stereoscope (Zeiss microscopy, Jena, Germany) equipped with a Canon EOS Rebel T3 digital camera (Canon, Tokyo, Japan).

### Live imaging of N. vectensis stained with Acridine Orange

*N. vectensis* embryos, polyps, and adults were maintained in 6-well plates filled to half depth with 1/3 ASW. Stained samples were maintained at room temperature during the initial hours of development and placed in 16 °C incubators for later stages of development and longer periods of husbandry. Samples were visualized in their husbandry plates under dark-field and florescence microscopy using a ZEISS Axio Zoom V16 stereoscope (Zeiss microscopy, Jena, Germany) equipped with filter sets for green (470/525 nm excitation/emission) and red (572/629) fluorescence and a Canon EOS Rebel T3 digital camera (Canon, Tokyo, Japan).

## RESULTS AND DISCUSSION

To establish and evaluate the effectiveness of AO as a tool for labeling *N. vectensis*, embryos were incubated in 1/3 ASW containing 0.664 µg/mL AO. Samples were left for 45 minutes in the absence (negative control) or presence of AO and washed five to seven times with 1/3 ASW to remove AO from husbandry media. The samples were then monitored throughout development under dark-field (Figure 1, A and B) and fluorescence microscopy (Figure 1, A’ and B’). The overall anatomy of polyps in the negative control group (Figure 1A) was indistinguishable to those exposed to AO (Figure 1B) when visualized under dark-field microscopy. However, when animals were visualized under green fluorescence microscopy, a clear difference was observed between autofluorescence in unstained animals (Figure 1A’) and labeling of tissue by incubation in AO (Figure 1B’). This initial experiment showed that despite some autofluorescence generated by *N. vectensis* (as reported by Ikmi and Gibson, 2010), AO can be used to enhance live visualization of this organism under fluorescence microscopy.

To establish an optimal concentration of AO that would allow for cellular labeling during embryogenesis without toxic outcomes, 10-fold dilutions of the dye were tested. Embryos were exposed for 45 minutes to 0.0664 µg/mL, 0.664 µg/mL and 6.64 µg/mL AO in 1/3X ASW, or 1/3X ASW alone, and both fluorescence intensity and survivability were assessed throughout development and metamorphosis. The survivability assay indicated statistically significant differences in mortality rates (*p* value < 0.05, ANOVA with Tukey’s Range Test) across all tested concentrations at at least two timepoints, suggesting that AO at these dosages does carry some level of toxicity to the embryos (Figure 1C). Embryos exposed to 0.0664 µg/mL and 0.664 µg/mL reached full development close to 70% of the time, which was close to the 80% success observed for negative controls raised in parallel (Figure 1C). In contrast, about 50% of embryos exposed to AO at a concentration of 6.64 µg/mL died by 30 days post-fertilization (dpf; Figure 1C). AO concentration at higher (66.4 µg/mL) and lower (0.00664 µg/mL) rates were also tested (data not shown) but were found not applicable because of almost complete lethality or very little signal detection, respectively. Consequently, the intermediate concentration of 0.664 µg/mL, which provided the best balance between labeling strength and survivability, was selected for subsequent experiments.

Lineage tracing by fluorescence labeling was attempted by microinjecting single *N. vectensis* blastomeres at the 4-cell stage with AO at a concentration of 66.4 µg/mL. This concentration was chosen as we calculated that the injected volume would be approximately 1/50 to 1/100 the cytoplasmic volume of each cell. Surprisingly, injected blastomeres did not retain fluorescent signal for longer than two minutes (Figure 2). In fact, signal from the dye dissipated quickly after injection, with the vast majority lost within 20 seconds (Figure 2, third panel), suggesting that AO is not applicable for cytoplasmic microinjection and that the cell surface is most likely the cellular content retaining AO upon immersion in ASW containing the dye during early stages of development.

**Figure 2.**
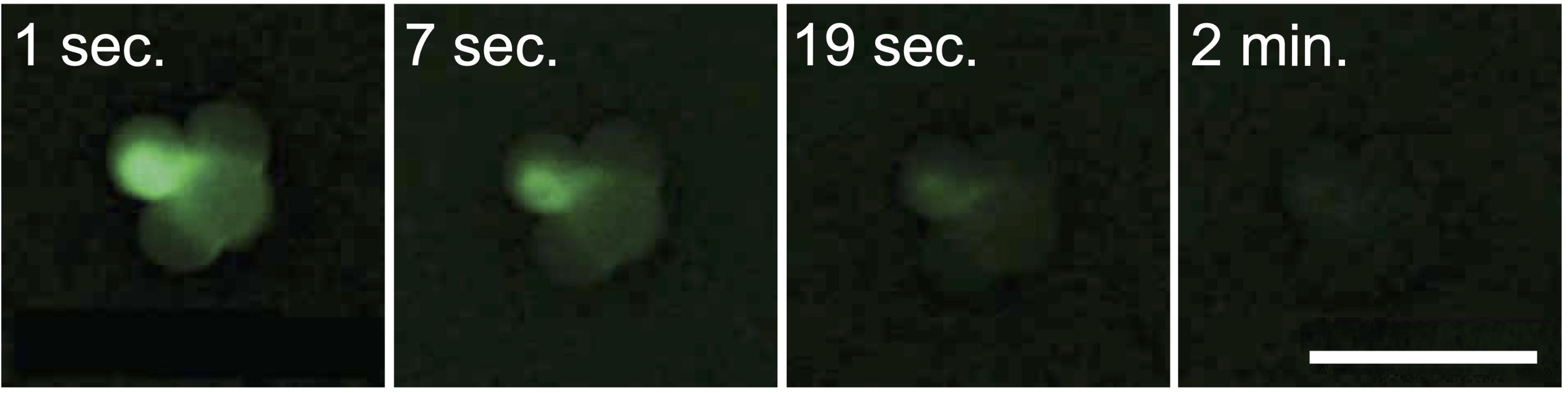
AO signal dissipates quickly upon injection in *N. vectensis* blastomeres. Time-lapse images show dissipation of AO green fluorescence signal following microinjection into a single blastomere from a 4-cell stage *N. vectensis* embryo. Time-points shown from left to right include 1 second, 7 seconds, 19 seconds, and 2 minutes following injection. Scale bar = 0.1 mm.

The longevity of the dye was tracked throughout development of *N. vectensis* by exposing zygotes to a single pulse of the dye by immersion into 1/3X ASW containing AO at a final concentration of 0.664 µg/mL, and imaging samples under darkfield, as well as green and red fluorescence microscopy (Figure 3). Initially, both green and red fluorescence signals were robustly detected along the surface of all cells in stained embryos, and the detection of these signals lasted for the first seven days of development (Figure 3, A-A” to I-I”). Interestingly, the red channel signal vanished after about eleven days and was hardly detectable at 31 dpf (Figure 3J”). On the other hand, the green signal was retained past 46 days, at which point enrichment was observed in specific subsets of cells. Cells labeled by fluorescence a month after exposure to AO were particularly visible as puncta in the tentacle region and down the body column as far as 46 and even 103 dpf (Figures 3, K’ and L’; Supplementary Video S1).

**Figure 3.**
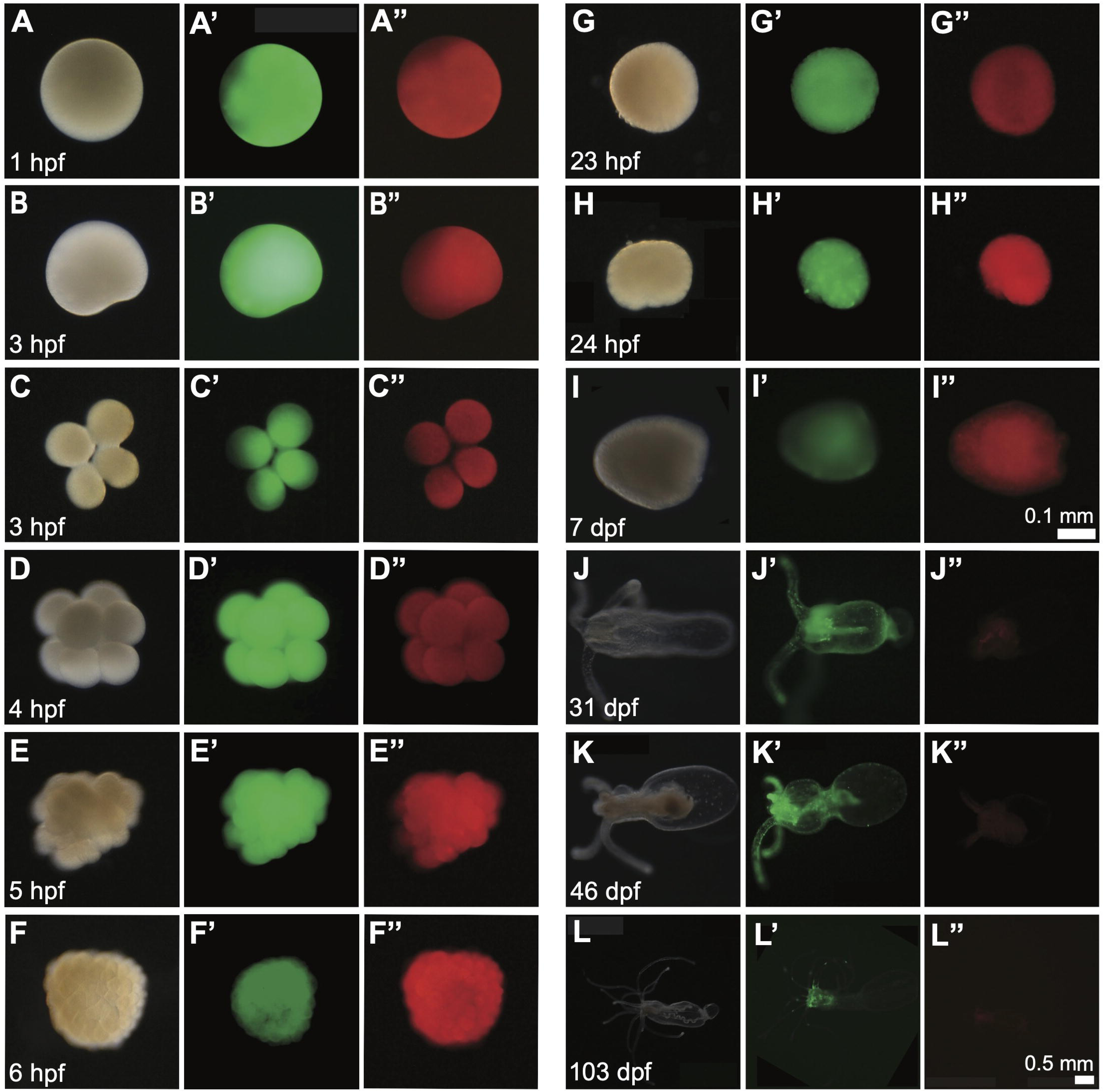
Exposure of *N. vectensis* zygotes to AO results in fluorescent labeling retained throughout development and metamorphosis. Dark-field (A-L), green florescence (A’-L’), and red florescence (A”-L”) microscopy images of live *N. vectensis* exposed to a 45-minute incubation in 0.664 µg/mL AO and visualized at one-hour post-fertilization (1 hpf; A-A”) and subsequent time-points (B-B” to K-K”) up to 103 days post-fertilization (103 dpf; L-L”). Scale bar for (A-I) is shown in (I; scale bar = 0.1 mm) and for (J-L) is shown in (L; scale bar = 0.5 mm)

To test for retention and distribution of AO during the process of regeneration, stained male polyps of more than a year in age underwent transverse amputations and analyzed for regeneration of the oral end (Figure 4A). One-hour post-amputation, green signal from AO was observed throughout the physa and remaining body column, which remained retracted following the injury (Figure 4B). Four and seven days following amputation, fluorescent green signal was visible throughout the pre-existing anatomy, but accumulation was observed in the regenerating end (Figure 4, C and D). By 17 days post-amputation, the oral was fully regenerated (Figure 4E). AO signal at this point was most intense in the mesenteries and tentacles but still present throughout the body column and physa (Figure 4E). By 41 days post amputation, the fluorescence from AO labeling could still be detected throughout the regenerated animal and mainly in the tentacles (Figure 4F). These observations indicate that AO is retained throughout regeneration and inherited by newly grown tissue.

**Figure 4.**
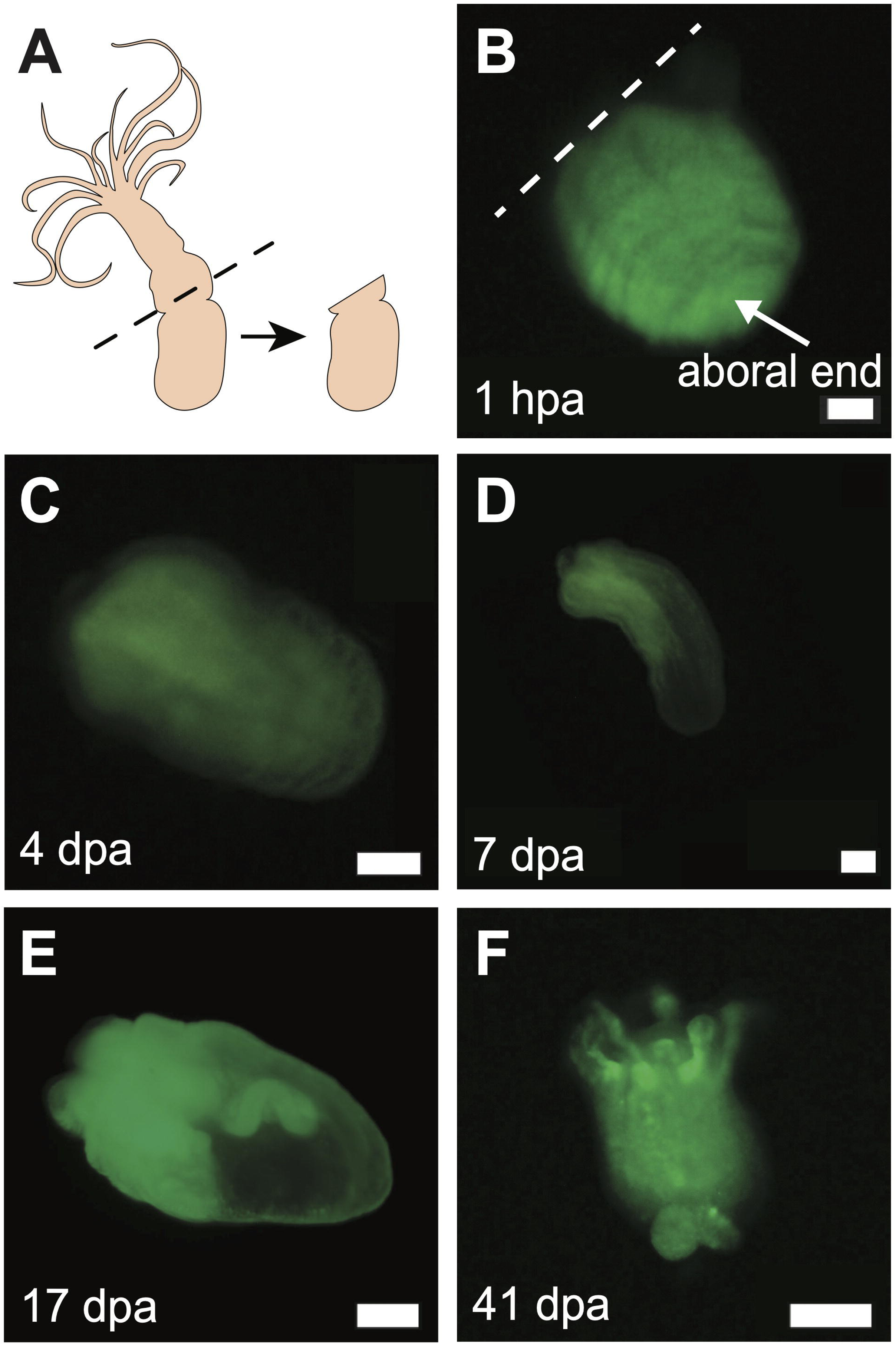
AO signal is from pre-labeled aboral ends is present in newly developed structures following regeneration. **(A)** Schematic shows transverse amputation of *N. vectensis* and aboral end analyzed during regeneration. **(B-F)** Images under green fluorescence microscopy show the aboral end of an AO-labeled polyp progressing through regeneration after transverse amputation. Images of the same sample were taken over the course of a month starting with 1-hour post-amputation (1 hpa; B), 4 days post-amputation (4 dpa; C), 7 dpa (D), 17 dpa (E), and 41 dpa (F). Scale bars = 0.1 mm.

Given the evidence for long-term retention of AO in *N. vectensis* and its utility for live imaging, we wondered whether it could be similarly used to visualize development of aquatic vertebrate models that undergo external development. To test this, zebrafish zygotes were stained with AO using the same concentration as described for *N. vectensis* and visualized under dark field and florescence microscopy at different timepoints during the first five days of development. AO green fluorescence labeling was detected in zebrafish blastomeres at 5 hours post fertilization (5 hpf), indicating effective initial uptake of the dye (Figure 5, A and A’). As development progressed, and as early as 2-days post-fertilization, the signal from most of the body of the embryo declined and accumulation was observed only in the yolk sack (Figure 5, C-D’), where it remained at 5-days post-fertilization (Figure 5, E-F’). Compared to *N. vectensis*, there was markedly reduced signal intensity in tissues of zebrafish embryos at within less than a week of staining.

**Figure 5.**
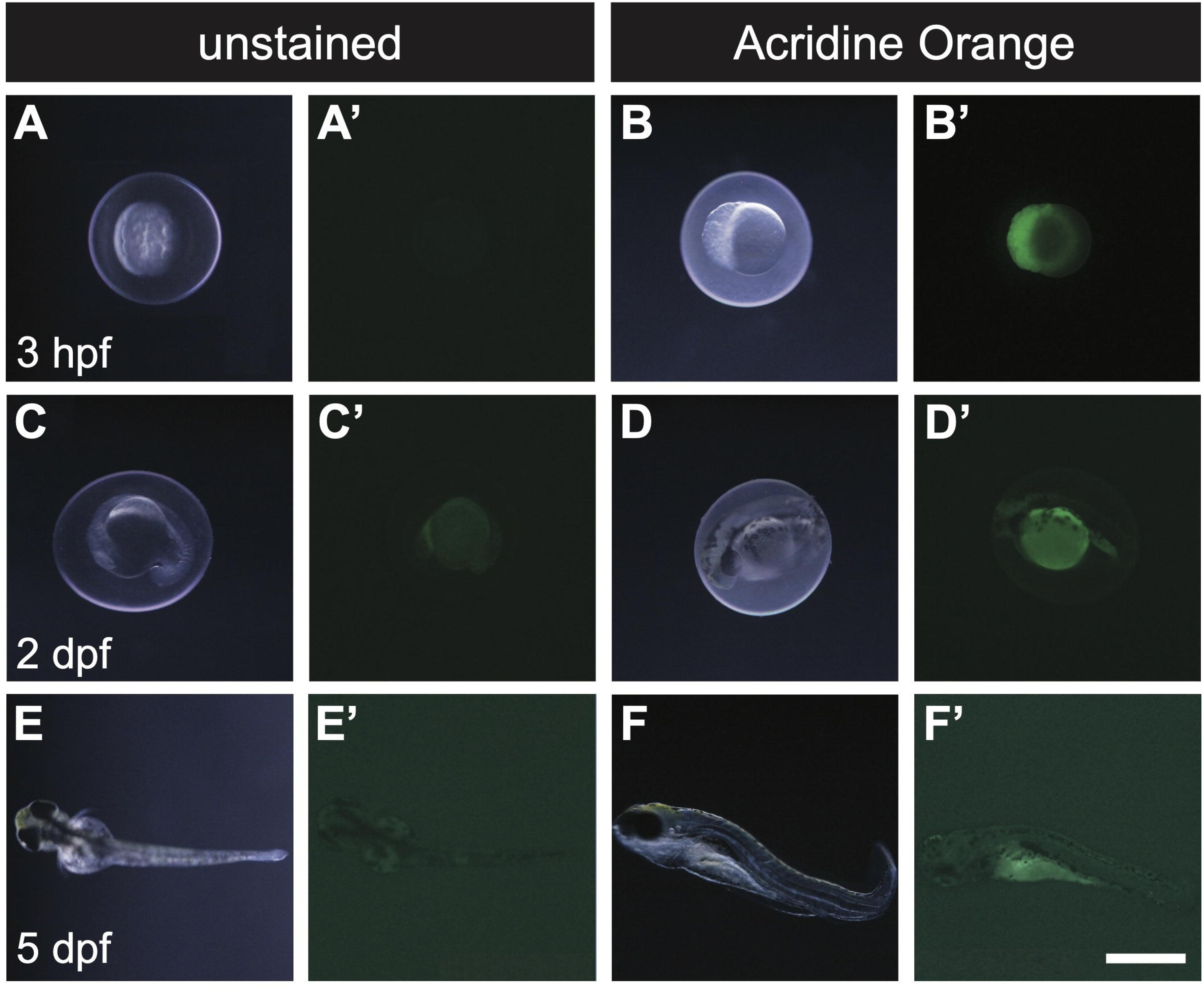
Brief retention of AO label in zebrafish embryos. **(A-F)** Darkfield (A-E) and florescence (A’-E’) live microscopy images of unstained (A-A’, C-C’, and E-E’) and AO-stained (B-B’, D-D’, and F-F’) zebrafish embryos and larvae . Green fluorescence is detected in AO-labeled blastomeres at 3 hpf (B’) and in the yolk sac at 2 dpf (D’) and 5 dpf (F’). Scale bar = 0.5 mm.

Our results demonstrate that a single, short-term exposure to Acridine Orange enables long-term fluorescence cellular labeling in *Nematostella vectensis* during development, homeostasis, and regeneration. This method provides a powerful tool for live imaging in a key early diverging metazoan. The applicability of AO for live imaging across animal phyla may be limited by differences in retention of dye signal, as observed in zebrafish (Figure 5) and planarian flatworms (data not shown), in which fluorescence signal is lost in less than a week. Analyses in other cnidarians, additional marine animals, as well as species belonging to even earlier-branching phyla (*e.g.*, sponges, ctenophores, and Placozoa) will reveal the breath of utility of AO as a long-term dye for fluorescence live imaging.

Important questions remain regarding the molecular basis of AO retention in *N. vectensis*, such as its use in cell and tissue transplantation experiments. Signal from AO labeling in *N. vectensis* does not seem to depend directly on DNA, RNA, or apoptotic cells (as described in mammals), so it is important to address what cellular entity or entities are labeled by AO. The observation of retention in the yolk sac of zebrafish suggests that labeling during early development in *N. vectensis* may be related to yolk protein. However, AO signal dissipated almost immediately when injected into blastomeres (Figure 2), indicating that perhaps AO is retained by entities in the cell surface and not in the cytoplasm. Additionally, as polyps grew into adulthood, the signal became concentrated in subsets of cells around the tentacles and pharynx, as well as in motile cells in the body column (Figure 3, K’ and L’; Supplementary Video S1). Presumably, cells in adults retain the AO label by different mechanisms than cells in the early embryo. Identifying the molecular entity (or entities) retaining AO may also help reveal the reason behind different red and green emission signal dynamics observed across *Nematostella vectensis* life stages.

An additional application for AO given its long-term, non-toxic retention in cnidarians, would be its use in field-based studies such as those in the context of marine conservation and reef restoration efforts. The capacity to non-invasively mark anemones, embryos, and polyps with AO provides a non-transgenic avenue to monitor survival, growth rates, and dispersal of individuals in different ecosystems and/or under different treatments. Monitoring labeled individuals in a heterogeneous population would provide critical data for evaluating the success of different restoration techniques for cnidarian species and their survival under different conditions.

## ACKNOWLEDGEMENTS

The authors thank Jessica MacNeil and Kavita Venkataramani from the Siegfried Lab at the University of Massachusetts Boston for their assistance with zebrafish larvae, as well as Nick Rivas for contributions during initial stages of this project. VP was supported by the Ralph and Janice Hames Family Foundation through the Professional Apprenticeship Career Experience Program (PACE) at the University of Massachusetts Boston. This work was supported by startup funds from the University of Massachusetts Boston to LR.

## REFERENCES

Amiel, A. R., Johnston, H. T., Nedoncelle, K., Warner, J. F., Ferreira, S., & Röttinger, E. (2015). Characterization of Morphological and Cellular Events Underlying Oral Regeneration in the Sea Anemone, *Nematostella vectensis*. International journal of molecular sciences, 16(12), 28449–28471.

Boero, F., Bouillon, J., & Piraino, S. (2005). The role of Cnidaria in evolution and ecology. Italian Journal of Zoology, 72(1), 65–71.

Burton, P. M., & Finnerty, J. R. (2009). Conserved and novel gene expression between regeneration and asexual fission in *Nematostella vectensis*. Development genes and evolution, 219(2), 79–87.

Carvalho, J. E., Burtin, M., Detournay, O., Amiel, A. R., & Röttinger, E. (2025). Optimized husbandry and targeted gene-editing for the cnidarian *Nematostella vectensis*. Development, 152(2). 10.1242/dev.204387

Chen C. Y., McKinney S. A., Ellington, L. R., & Gibson, M. C. (2020). Hedgehog signaling is required for endomesodermal patterning and germ cell development in the sea anemone *Nematostella vectensis*. Elife, 9:e54573.

Cheung, S., Bredikhin, D., Gerber, T., Steenbergen, P. J., Basu, S., Bailleul, R., Hansen, P., Paix, A., Benton, M. A., Korswagen, H. C., Arendt, D., Stegle, O., & Ikmi, A. (2025). Systemic coordination of whole-body tissue remodeling during local regeneration in sea anemones. Developmental cell, 60(5), 780–793.e7.

Denner, A., Steger, J., Ries, A., Morozova-Link, E., Ritter, J., Haas, F., Cole, A.G., & Technau, U. *Nanos2* marks precursors of somatic lineages and is required for germline formation in the sea anemone *Nematostella vectensis*. Sci Adv. 10(33):eado0424. doi: 10.1126/sciadv.ado0424.

Doyle, T.K., Hays, G.C., Harrod, C., & Houghton, J.D.R. (2014). Ecological and Societal Benefits of Jellyfish. In: Pitt, K., Lucas, C. (eds) Jellyfish Blooms. Springer, Dordrecht.

DuBuc, T. Q., Traylor-Knowles, N., & Martindale, M. Q. (2014). Initiating a regenerative response; cellular and molecular features of wound healing in the cnidarian *Nematostella vectensis*. BMC biology, 12, 24

DuBuc, T. Q., Dattoli, A. A., Babonis, L. S., Salinas-Saavedra, M., Röttinger, E., Martindale, M.Q., & Postma M. (2014). In vivo imaging of *Nematostella vectensis* embryogenesis and late development using fluorescent probes. BMC Cell Biol., 15:44. doi: 10.1186/s12860-014-0044-2.

Ettinger, A., & Wittmann, T. (2014). Fluorescence live cell imaging. Methods in cell biology, 123, 77–94.

Finnerty, J.R., Pang, K., Burton, P., Paulson, D., & Martindale, M. Q. (2004). Origins of bilateral symmetry: *Hox* and *dpp* expression in a sea anemone. Science, 304(5675):1335–7. doi: 10.1126/science.1091946.

Finnerty, J. R., Paulson, D., Burton, P., Pang, K., & Martindale, M. Q. (2003). Early evolution of a homeobox gene: the parahox gene Gsx in the Cnidaria and the Bilateria. Evolution & Development, 5(4), 331–345.

Galliot, B. (2012). Hydra, a fruitful model system for 270 years. The International journal of developmental biology, 56(6-8), 411–423. 10.1387/ijdb.120086bg

Hand, C., & Uhlinger, K. R. (1992). The culture, sexual and asexual reproduction, and growth of the sea anemone *Nematostella vectensis*. Biological Bulletin, 182(2), 169–176.

Ikmi, A. & Gibson, M. C. (2010). Identification and in vivo characterization of NvFP-7R, a developmentally regulated red fluorescent protein of *Nematostella vectensis*. PLoS One. 5(7):e11807.

Layden, M. J., Rentzsch, F., & Röttinger, E. (2016). The rise of the starlet sea anemone *Nematostella vectensis* as a model system to investigate development and regeneration. Wiley Interdiscip Rev Dev Biol, 5(4):408–28.

Layden, M. J., Röttinger, E., Wolenski, F. S., Gilmore, T. D., & Martindale, M. Q. (2013). Microinjection of mRNA or morpholinos for reverse genetic analysis in the starlet sea anemone, *Nematostella vectensis*. Nature protocols, 8(5), 924–934.

Martindale, M. Q., Pang, K., & Finnerty, J. R. (2004). Investigating the origins of triploblasty: “Mesodermal” gene expression in a diploblastic animal, the sea anemone *Nematostella vectensis* (Phylum Cnidaria; Class Anthozoa). Development, 131(10), 2463–2474.

Miramón-Puértolas, P., Pascual-Carreras, E. & Steinmetz, P.R.H. (2024). A population of Vasa2 and Piwi1 expressing cells generates germ cells and neurons in a sea anemone. Nat Commun 15, 8765.

Paix, A., Basu, S., Steenbergen, P., Singh, R., Prevedel, R., & Ikmi, A. (2023) Endogenous tagging of multiple cellular components in the sea anemone *Nematostella vectensis*. Proc Natl Acad Sci. 3;120(1):e2215958120.

Passamaneck, Y. J., & Martindale, M. Q. (2012). Cell proliferation is necessary for the regeneration of oral structures in the anthozoan cnidarian *Nematostella vectensis*. BMC Developmental Biology, 12, 34.

Plemel, J. R., Caprariello, A. V., Keough, M. B., Henry, T. J., Tsutsui, S., Chu, T. H., Schenk, G. J., Klaver, R., Yong, V. W., & Stys, P. K. (2017). Unique spectral signatures of the nucleic acid dye acridine orange can distinguish cell death by apoptosis and necroptosis. Journal of Cell Biology, 216(4), 1163–1181.

Rossi, S., Gravili, C., Milisenda, G., Bosch-Belmar, M., De Vito, D., & Piraino, S. (2019). Effects of global warming on reproduction and potential dispersal of Mediterranean Cnidarians. The European Zoological Journal, 86(S1), 255–271.

Ryan, J. F., Mazza, M. E., Pang, K., Matus, D. Q., Baxevanis, A. D., Martindale, M. Q., & Finnerty, J. R. (2007). Pre-Bilaterian Origins of the Hox Cluster and the Hox Code: Evidence from the Sea Anemone, *Nematostella vectensis*. PLoS ONE, 2(1), e153.

Stefanik, D.J., Friedman, L. E, & Finnerty, J. R. (2013). Collecting, rearing, spawning and inducing regeneration of the starlet sea anemone, *Nematostella vectensis*. Nature Protocols, 8(5):916–923.

Trembley, A. (1744). Mémoires pour servir à l’histoire d’un genre de polypes d’eau douce, à bras en forme de cornes. Par A. Trembley. Chez Jean & Herman Verbeek.

Warner, J. F., Guerlais, V., Amiel, A. R., Johnston, H., Nedoncelle, K., & Röttinger, E. (2018). NvERTx: A gene expression database to compare embryogenesis and regeneration in the sea anemone *Nematostella vectensis*. Development, 145(10), dev162867.

Weis, V. M. (2008). Cellular mechanisms of Cnidarian bleaching: stress causes the collapse of symbiosis. Journal of Experimental Biology, 211(19), 3059–3066.

