## Supplementary Video S1 legend for "Acridine Orange dye for long-term staining and live imaging of cnidarian development and regeneration"

**SUPPLEMENTARY MATERIAL**

**Supplementary Video S1. Live imaging of Acridine Orange-labeled *N. vectensis* polyp under green florescence microscopy.** *N. vectensis* polyps recorded under green fluorescence microscopy (A’ and B’) shows retention of AO label obtained during a 45-minute incubation at the zygotic stage of development. Enrichment of AO signal is detected in cells of the tentacles and gastrovascular cavity.
